# Interspecific variation of olfactory preferences in flies, mice, and humans

**DOI:** 10.1101/358713

**Authors:** Diogo Manoel, Melanie Makhlouf, Antonio Scialdone, Luis R. Saraiva

## Abstract

Aiming to unravel interspecific differences in olfactory preferences, we performed comparative studies of odor valence in flies, mice, and humans. Our analysis revealed that odor preferences of flies correlate positively with the ones of mice and negatively with the ones of humans, but found no evidence supporting the hypothesis that humans and mice prefer the same odors. We further find that odorants eliciting the highest and lowest preferences are often advertising critical biological sources (e.g., food or oviposition sites), suggesting that evolutionary pressures reflecting the ecological needs of each species shape olfactory preferences.

---

Animals perceive myriad odors as pleasant or neutral and yet others as unpleasant or repugnant. Olfactory cues can also elicit changes in behavior and physiology, thus playing an instrumental role in survival, reproduction and species-specific adaptations to different ecological niches^1,2^. Of the many roles of olfaction, it has recently been proposed that assessing the valence of odors is its key function in humans^3^.However, such perceptions and rating of odorants across a hedonic scale can be innate, learned, and modulated by the internal state of the individual, or even be context dependent^1,3,4^. Moreover, while many molecular mechanisms underlying olfaction are conserved between species, rapid evolutionary dynamics ensure the creation of highly species-specific repertoires and relative abundances of olfactory receptors (ORs), which ultimately shape the olfactory preferences and abilities in different animals^2,5^. Under this assumption, it would be fair to assume that the higher the evolutionary distance between two given species, the more their olfactory preferences would diverge. Exceptions to this assumption could exist for small groups of ecologically relevant odors common between two given species, or even for distantly related species with converging biology or ecological niches.

Recent studies comparing olfactory preferences between flies-humans and mouse-humans revealed that the perception of odor intensity and pleasantness are conserved in these species pairs, respectively, and that judgments of odor quality are different in fruit flies-humans^6,7^. Taken together, these observations suggest that distinct aspects of olfactory perception can be either species-specific or conserved across these evolutionarily distant species pairs. However, these prior studies used small sample sizes and different criteria to select the odorants, which could impact the interpretations of these findings. This is mainly due to highly combinatorial nature of the olfactory systems, allied to the vast arrays of odorants present in nature and the large receptor repertoires equipping individual species^1,8–10^. Fortuitously, the odor valences (i.e., olfactory preferences) of large panels of odorants (73-480) have recently been characterized in flies (*Drosophila melanogaster*), mice (*Mus musculus*) and humans (*Homo sapiens*)^4,11,12^. Using these data as a starting point, we performed comparative studies aimed at dissecting the differences in olfactory preferences among these three species.

We started by compiling the odor valence scores for all possible combinations of overlapping odorants between flies, mice, and humans (Fig. 1a). Our first observation was that, for both flies and mice, odorants associated with putrefaction^13^ and fermentation^14–16^ (e.g., propionic acid, putrescine, 2-phenylethanol) are among the most attractive odorants, while odorants depleted in non-decaying fruits^17,18^(e.g., benzaldehyde, linalool, octanol) elicit opposite behaviors (Supplementary File 1). Next, we compared the attraction indexes (AI) of flies and the olfactory investigation times (OIT) of mice for the 16 overlapping odorants and found a strong positive correlation (rs=0.700, p=0.003, Fig. 1b and Supplementary File 2) between the olfactory preferences of these species. These findings are consistent with the ecological habits of flies and mice, as both use odors emanating from feces, fermenting or decaying organic material (such as fruits or meat) as cues for essential aspects of their biology, such as feeding or oviposition^1,10,19–25^.

**Figure 1.**
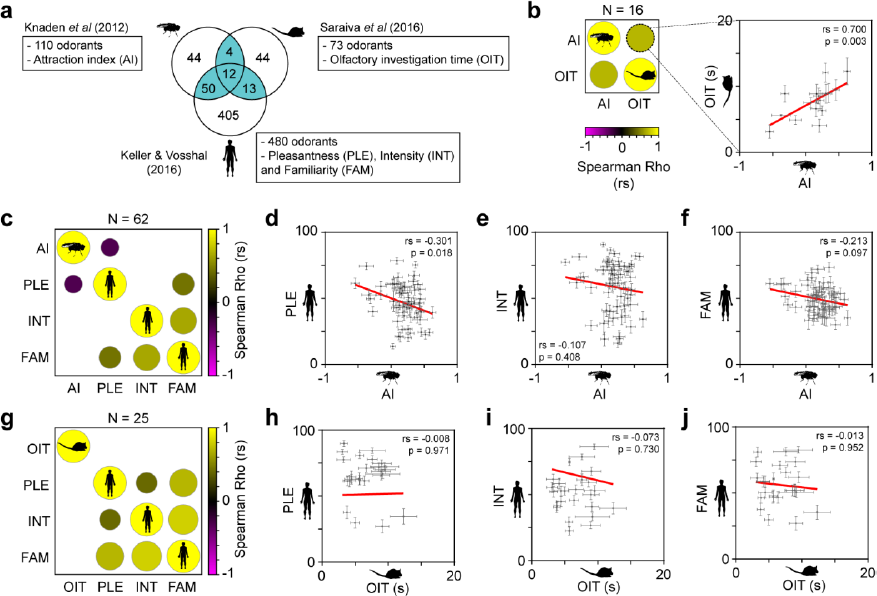
Interspecific differences of odor valence in fruit flies, mice and humans. **a** – Study design: the Venn diagram indicates the number of odorants (highlighted in ocean blue) for which odor valence scores overlap in flies^11^, mice^4^, and humans^12^. For each species, the name and abbreviation of the parameters measured are specified as follows: fly attraction index (AI), mouse olfactory investigation time (OIT) and human-rated pleasantness (PLE), intensity (INT) and familiarity (FAM). **b** – Correlogram matrix (left) and correlation plot (right) comparing the fly AI with mouse OIT for the 16 overlapping odorants. **c** - Correlogram for the 62 overlapping odorants between fly AI and the human PLE, INT and FAM. To the right are the corresponding correlation plots between fly AI and human-rated PLE (**d**), INT (**e**) and FAM (**f**). **g** - Correlogram for the 25 overlapping odorants between mice OIT and humans PLE, INT and FAM. To the right are the corresponding correlation plots between mouse OIT and human-rated PLE (**h**), INT (**i**) and FAM (**j**). In all correlograms, only significant (p<0.05) correlations are plotted, and the circle size and color indicate the magnitude and direction of the correlation (Spearman rho, rs). Blank cells correspond to non-significant correlations.

We then focused on the set of 62 intersecting odorants between flies and humans, and compared the AI of flies to three measures of human odor perception: pleasantness (PLE), intensity (INT) and familiarity (FAM) (Fig. 1c-f). We found that AI of flies correlates negatively with the human-rated PLE (rs=-0.301, p=0.018, Fig. 1d and Supplementary File 2), but not with INT or FAM (Fig. 1e, f). We observed that the human odor valence parameters tend to correlate positively with each other (Fig. 1c, g, Supplementary Fig. 1b and Supplementary File 2), in line with previous studies^12,26^. As mentioned above, among the most attractive odorants to flies are molecules associated with putrefaction^13^ and fermentation^14–16^ (e.g., isobutyric acid, propionic acid, 3-(methylthio)-1-propanol), which are unpleasant to humans (Supplementary File 1). In contrast, odorants associated with ripening fruits^17,18^ (e.g., benzaldehyde, linalool, octanol) score high in the human pleasantness rating but are aversive to flies. Once again, these findings are consistent with the feeding ecology of these species, as humans favor fruits at different stages or ripening and flies preferentially feed on decaying or fermenting fruits ^22^.

Subsequently, we tested the odor valence scores for the intersecting 25 odorants between the mouse and human studies. Here, we found no significant correlation between the mouse OIT and the human-rated PLE, INT, or FAM (Fig. 1g-j and Supplementary File 2). Again we observe that odorants associated with putrefaction^13^ and alcoholic fermentation processes^27^ (e.g., propionic acid, pyrrolidine), are attractive to mice, but unpleasant to humans (Supplementary File 1). These results are consistent with the hypothesis that mice and humans use these cues to seek and avoid food, respectively, accordingly to their feeding habits. Interestingly, we noticed that odorants eliciting the highest pleasantness ratings in humans and aversive responses in mice are highly abundant in herbs, spices and medicinal oils (e.g. (+)-camphor, hexanol, 2-isobutylthiazole, (-)-fenchone). Such molecules are also known as plant secondary metabolites (PSMs), which, if ingested, can trigger a broad range of physiological effects, ranging from digestion impairment to toxicity^28,29^. Because mice are generalist feeders, their physiological ability to eliminate PSMs from the body is limited, and as a consequence, they might learn through physiological feedback to avoid toxic plants and their respective odor signatures^23,28–30^. Since humans are also generalists, what could explain the behavioral difference to PSMs between mice and humans? The most parsimonious explanation is perhaps that throughout evolution, the direct benefits of PSMs liking and consumption were able to offset the costs associated with the detoxification processes. Such benefits could derive from the antimicrobial, antifungal and healing properties of specific PSMs, which could explain the widespread human use of spices and plant essential oils for both gastronomic and medical uses^31,32^.

A limitation of the present study is the difference in the combinations of intersecting odorants used for the three-way interspecific comparisons. To address this, we performed an analysis using the odor valence ratings for all twelve overlapping odorants between the three studies (Supplementary Fig. 1a, b and Supplementary File 2). Consistent with the results above, we observe a strong positive correlation (rs=0.657, p=0.024) between flies and mice, but no correlation between mice and humans (Supplementary Fig. 1b). However, the negative correlation observed above between flies AI and rated PLE in humans (Fig. 1c) is no longer maintained, likely due to the ~5.5-fold reduction in odorant sample size.

Together, our data are consistent with a previous study showing that the judgments of odor quality between flies and humans are species-specific^6^. However, the present study contradicts the previously reported finding that a positive correlation exists between the olfactory preferences of mice and humans^7^. It is likely that this discrepancy arises from the different experimental protocols used to assess the mouse odor preferences, along with the dissimilar number, combinations and concentrations of odorants tested^4,7^.

In conclusion, our data support a model where flies and mice share similar olfactory preferences, but neither species share odor preferences with humans. These results are also consistent from an ecological perspective, as many odorants that elicit attraction in flies and mice, also serve as critical sensorial cues for several aspects of their biology^1,10,33^. In turn, such odorants are perceived as foul or unpleasant by humans, possibly functioning as a warning regarding the toxicity of their source. Future studies aimed at dissecting the olfactory preferences in a higher number of odorants, and across more species, are necessary to both stress-test the model proposed here and identify other associations.

## Methods

### Data source

Odor valence scores from flies (*Drosophila melanogaster*), mice (*Mus musculus*) and humans (*Homo sapiens*) were collected from Knaden *et al* (2012)^11^, Saraiva *et al* (2016)^4^ and Keller and Vosshall (2016)^12^, respectively. Odorants were presented at a dilution of 10^−1^-10^−3^ in flies^11^ and at a concentration of 85mM in mice^4^. For consistency purposes between the three studies, only the valence scores for the high concentrations in humans^12^ were used. Calculated data used to perform the analysis are available in the Supplementary File 1.

### Statistical analyse

Statistical analyses were done using GraphPad Prism (version 6.04), PAlaeontological STatistics (version 3.14)^34^ and the R statistical package^35^. Data points shown on all plots represent the averages and associated standard error of the mean (sem). Correlograms were generated using standard R functions *cor* and *cor.mtest* with Spearman method (95% confidence interval and α=0.05). Correlations matrices were visualized using *corrplot* method from the Bioconductor *corrplot* (v0.84) package^36^. We studied the significant association of the sample annotations (Supplementary Fig. 1B) to different groups by performing hierarchical clustering (HC) analysis on standardized data values and using Euclidean distances with Ward’s method.

### Data availability

Data used to perform the analysis are available (mean and sem) in Supplementary File 1. All Spearman’s correlation coefficients (rs) and associated p-values (P) calculated for all correlograms and correlation plots shown in this study are available in Supplementary File 2.

## Author Contributions

D.M. analyzed data and wrote the paper; M.M. analyzed data; A.S. contributed analysis tools, and L.R.S. conceived and supervised the project, analyzed data and wrote the paper.

## Acknowledgments

We would like to thank Bernice Lo for helpful comments and discussions, and Markus Knaden for providing us with the raw data reported in the Knaden *et al* (2012)^11^ study.

## Supplementary Material

**Supplementary Figure 1.**
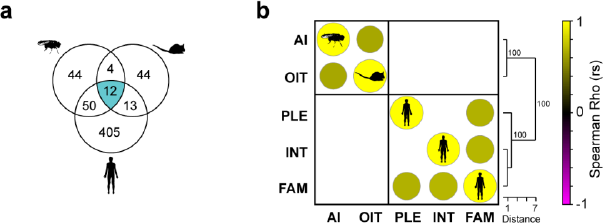
**Interspecific differences of odor valence in fruit flies, mice and humans** a - Venn diagram indicating the number of odorants (highlighted in ocean blue) for which odor valence scores overlap in all three species analyzed here. b - Correlogram comparing all the odor valence parameters quantified for the 12 overlapping odorants: fly attraction index (AI), mouse olfactory investigation time (OIT) and human-rated pleasantness (PLE), intensity (INT) and familiarity (FAM). Only significant correlations are plotted, and the circle size and color indicate the magnitude and direction of the correlation (Spearman’s rho, rs). Blank cells correspond to non-significant correlations. A hierarchical clustering analysis (to the right of the matrix) revealed the existence of 2 groups, one including the fly AI and mouse OIT, and the other composed by human odor valence parameters only. These two groups are supported by high bootstrap values (1-100).

### Supplementary File 1

Excel tables with the data taken from Knaden *et al* (2012)^11^, Saraiva *et al* (2016)^4^ and Keller and Vosshall (2016)^12^ used for the correlograms and plots.

### Supplementary File 2

Excel tables are containing all the Spearman’s correlation coefficients (rs) and associated p-values (P) calculated for all the correlograms shown in this study.

